# OGG1-Binding to Oxidized Guanine Base in Viral DNA Overcomes Epstein–Barr Virus Latency

**DOI:** 10.64898/2026.06.05.728896

**Authors:** Wenjing Hao, Xiaoqi Wang, Jiabo Li, Gang Liu, Lang Pan, Xueqing Ba, Istvan Boldogh, Ziyuan Duan

**Affiliations:** Institute of Genetics and Developmental Biology, Chinese Academy of Sciences, Beijing, China; Department of Infectious Diseases, Key Laboratory of Major Diseases in Children, Beijing Children’s Hospital, Ministry of Education, Capital Medical University, National Center for Children’s Health, Beijing, China; NHC Key Laboratory of Radiobiology, School of Public Health, Jilin University, Changchun, China; The Key Laboratory of Molecular Epigenetics of the Ministry of Education, School of Life Science, Northeast Normal University, Changchun, China; Department of Microbiology and Immunology, University of Texas Medical Branch at Galveston, Galveston, Texas, US

**Keywords:** EBV reactivation, lytic gene expression, ROS, oxidative base modification, OGG1 enrichment

## Abstract

Epstein–Barr virus (EBV) establishes lifelong latency in human cells and can periodically reactivate, contributing to several cancers. However, the molecular mechanisms that disrupt EBV latency, particularly those driven by oxidative stress, require further investigation. In this study, we provide evidence that oxidative DNA damage, particularly the formation of 8-oxoguanine and its repair enzyme OGG1, is involved in EBV reactivation. We demonstrated that OGG1 is recruited to EBV regulatory regions under oxidative stress conditions and is associated with the transcriptional activation of immediate-early and early lytic genes. Further experiments showed that pharmacological inhibition of OGG1 DNA binding significantly suppressed EBV lytic gene expression, supporting a functional role for OGG1 in viral reactivation. Mechanistically, our findings suggest that OGG1 may contribute to EBV lytic activation through a noncanonical mechanism independent of its glycosylase activity. These findings provide insight into how EBV may exploit host oxidative DNA damage responses to facilitate latency disruption and suggest that targeting OGG1 may offer a potential strategy to limit EBV reactivation and EBV-associated diseases.

**Graphical Abstract:** 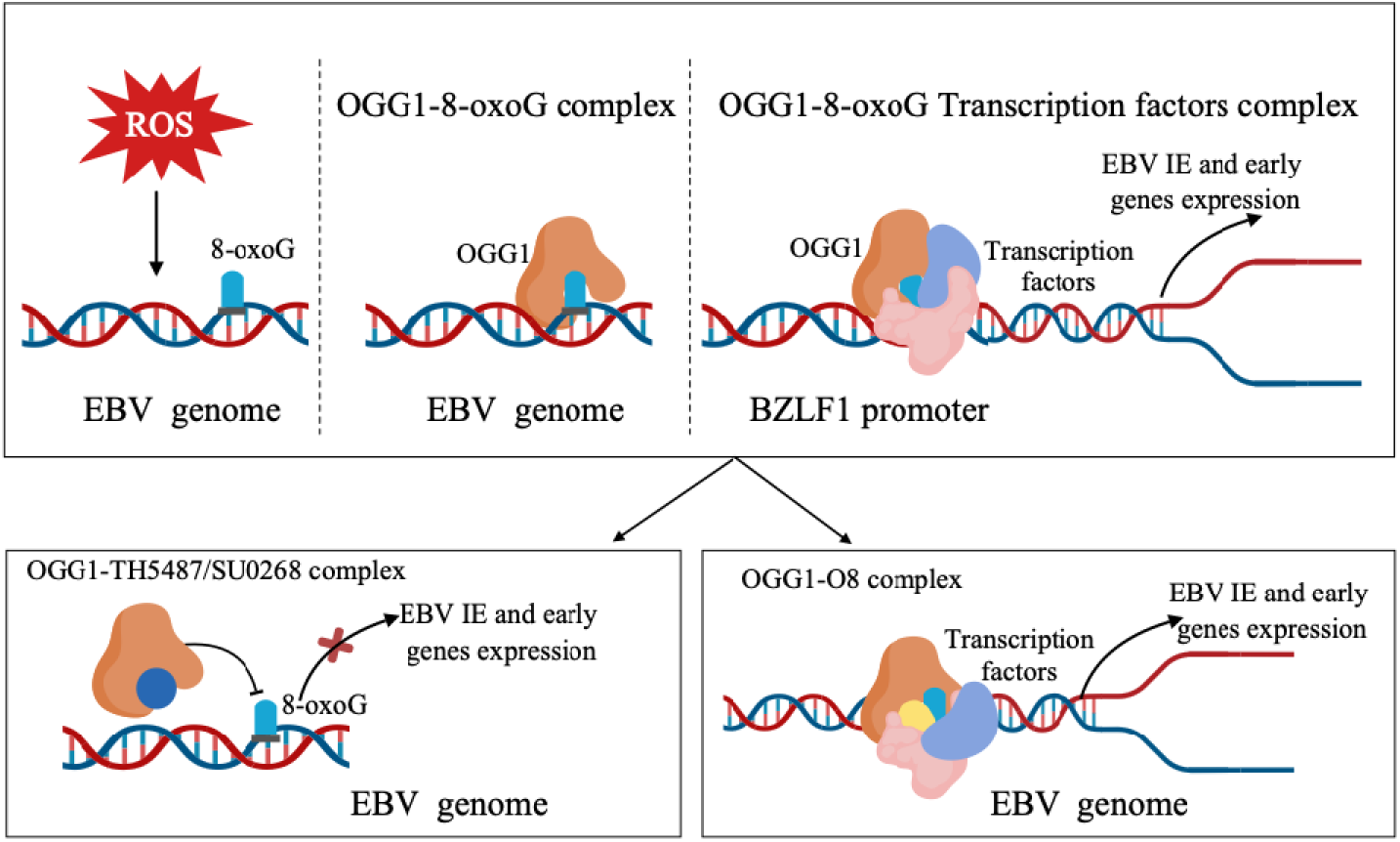

**The model illustrates how oxidative DNA damage promotes Epstein–Barr virus (EBV) lytic reactivation through host base excision repair machinery:** Reactive oxygen species (ROS) may induce oxidative modification of guanine to 8-oxoguanine (8-oxoGua) in latent EBV genomes, potentially contributing to OGG1 recruitment to viral regulatory regions. OGG1 binding facilitates the transcriptional activation of immediate-early and early lytic genes, likely through transcription factor recruitment, thereby contributing to the transition from latency to lytic replication. Pharmacological inhibition of OGG1-DNA interaction suppresses this process, highlighting OGG1 as a potential therapeutic target. 8-oxoGua, 7,8-dihydro-8-oxoguanine; OGG1, 8-oxoguanine DNA glycosylase-1.

**Significance Statement:**

- identifies a potential mechanism by which ROS-induced oxidative DNA base damage and OGG1 recruitment within the EBV genome may contribute to viral reactivation from latency.
- suggests that oxidative DNA repair intermediates may serve as regulatory platforms for viral gene expression.
- provides pharmacological evidence supporting OGG1 as a potential therapeutic target for suppressing EBV reactivation.

## 1. Introduction

Epstein-Barr virus (EBV) infects up to 95% of the human population, with most infections remaining asymptomatic. This is largely due to the ability of EBV to establish latency following a phase of limited progeny virion production. EBV can periodically reactivate into a lytic phase. Both primary and reactivated EBV infections are associated with human pathologies, ranging from benign conditions to malignant diseases (1, 2). EBV is a double-stranded DNA virus, and its genome size ranges from 170 to 184 kilobase pairs, encodes approximately 80 open reading frames (ORFs), expressing over 100 genes and ∼40 non-coding RNAs(3). These viral genes exhibit tightly regulated temporal expression patterns during latent and lytic phases(4).

Recent evidence indicates that both primary and reactivated EBV infections are accompanied by increased production of reactive oxygen species (ROS), which activate redox-sensitive signaling pathways, promote chromatin remodeling, and enhance the transcription of immediate-early and early lytic genes, thereby facilitating viral replication (5, 6). Notably, ROS are not only byproducts of infection but actively modulate the viral life cycle, shaping the balance between latency and lytic reactivation (7, 8). However, the mechanisms by which ROS promote EBV replication and reactivation from latency remain only partially understood.

ROS activates a complex network of signaling pathways, activating nuclear factor-kappa B (NF-κB), transcription factor (ATF)/c-JUN (c-FOS), cAMP response element-binding protein (CREB), interferon response factor 7 (IRF7), and the pro-oncogene c-MYC, known to be implicated in reactivation and the lytic cycle of EBV(5, 9). Beyond its role in signaling, ROS also induces oxidative modifications of cellular biomolecules, particularly nucleic acids. EBV is also known to hijack DNA damage response pathway to orchestrate primary infection and lytic reactivation(10). One of the predominant DNA base lesions is 7,8-dihydro- 8-oxoguanine (8-oxoGua), due to guanine’s lowest oxidation potential among nucleic acid bases and in nucleoside triphosphate pools(11, 12). In EBV-carrying immortalized B cells, enzymes including the nucleoside triphosphatase mut-T homolog 1 (MTH1), 8-oxoguanine DNA glycosylase-1 (OGG1) and adenine/guanine-specific DNA glycosylase (E. coli. Mut Y homolog) (13, 14).

In both viral and host DNA, 8-oxoGua is repaired via the base excision repair(BER) pathway, initiated by OGG1(15). Notably, the OGG1-8-oxoGua complex is not merely a DNA repair intermediate. It has been shown that this complex could function as a guanine nucleotide exchange factor (GEF) for the small GTPase RAS, promoting the conversion of RAS from the GDP-bound inactive state to the GTP-bound active state, thereby activating downstream RAS-MAPK and NF-κB signaling pathways involved in inflammatory and proliferative responses(16, 17). In addition, recent findings also show that 8-oxoGua in regulatory sequences is associated with altered CpG methylation patterns, functions as an epigenetic-like mark, and along with the bound OGG1 serves as a platform for binding of chromatin modifiers, trans-acting factors, and regulates gene expression(18–21).

Given roles of ROS in both primary and reactivation EBV infections, the redox susceptibility of intrahelical guanine and transcriptional coregulatory role of OGG1, we therefore hypothesized that OGG1 plays a key role in activation of the EBV lytic cycle. To test this, we utilized Akata-EBV, C666-1, and B95.8 cells, representing EBV latency types I, II, and III, respectively. Cells were co-treated with TPA and sodium butyrate (NaB) to induce EBV reactivation. Our results demonstrated that OGG1 binds to the 8-oxoGua within EBV promoter sequences and facilitates the expression of BZLF1 (Zta), consequently BALF2, and BMRF1 (EA-D) at both mRNA and protein levels, contributing to the transition from latency to productive viral replication. Importantly, inhibition of OGG1’s substrate binding but not its glycosylase activity, suppressed EBV reactivation. Taken together, these results suggest that ROS-induced Gua modification and OGG1 binding act as a molecular switch controlling the transition from latency to lytic EBV replication.

## 2. Materials and Methods

### 2.1 Reagents

The polyclonal Rabbit antibody (Ab) against OGG1 (Cat# NB100-106) was purchased from Novus Biologicals (Colorado, USA). The monoclonal antibody (Ab) against EBV ZEBRA (Zta, Cat# sc-53904) and EBV Ea-D (BMRF1, Cat# sc-58121) was purchased from Santa Cruz (Texas, USA). The monoclonal antibody against GAPDH (Cat# 60004), P38 (Cat# 14064-1-AP), JNK (Cat# 24164-1-AP), ERK (Cat# 16443-1-AP), Phospho-p38 MAPK (Thr180/Tyr182) (Cat# 28796-1-AP), Phospho-JNK (Tyr185) (Cat# 80024-1-RR) and Phospho-ERK1/2 (Thr202/Tyr204) (Cat# 28733-1-AP) antibody were purchased from Proteintech (Wuhan, Hubei, China). Phorbol 12-myristate 13-acetate (TPA, P8139) and Sodium butyrate (NaB, 303410) were purchased from Sigma (Saint Louis, MO, USA). The secondary antibody HRP-Goat Anti-Mouse IgG (SA00001-1), HRP-Goat Anti-Rabbit IgG (SA00001-2) were purchased from Proteintech (Wuhan, Hubei, China). OGG1 inhibitor, TH5487 (4-(4-Bromo-2-oxo-3H-benzimidazol-1-yl)-N-(4-iodophenyl) piperidine-1-carboxamide) (Cat# HY-125276) and SU0268 (modified acyl tetrahydroquinoline sulfonamide skeleton) (Cat# HY-139056) was purchased from MedChem Express (Shanghai, China). O8 (3,4-dichloro-benzo[b]thiophene-2-carboxylic acid hydrazide) (Cat# SML1697) was purchased from Sigma (Saint Louis, MO, USA). All inhibitors were dissolved in DMSO, and equivalent vehicle controls were included in all experiments. The final DMSO concentration was kept consistent across all treatment groups.

### 2.2 Cell culture and treatment

Akata-EBV and B95.8 cells were grown in RPMI 1640 (Gibco, Thermo Fisher Scientific, USA) supplemented with 10% of fetal bovine serum (Procell Cat# 164210, Wuhan, Hubei, China) and 100 IU penicillin and 100 µg streptomycin at 37°C in a 5% CO2 atmosphere. C666-1 cells (Misson Cell Biotechnology, Cat# CTCC-400-0109) were cultured in RPMI 1640 (Misson Cell Biotechnology, Cat# CTCC-002-003) and 10% fetal bovine serum (Misson Cell Biotechnology, Cat# CTCC-002-071) (Zhejiang, China). Akata-EBV cells were kindly provided by Tao Peng (Sino-French Hoffmann Institute, Guangzhou Medical University, Guangzhou, Guangdong, China.). Cells were treated with solvent (mock) or treated with 10 µM OGG1 inhibitor TH5487, O8 or SU0268 for one hour respectively, followed by treatment with 20 ng/ml TPA and 3mM NaB for various lengths of time. The concentration of 10 µM was selected based on previous reports demonstrating effective inhibition of OGG1 activity without significant cytotoxicity.

### 2.3 Quantitative reverse transcription real-time PCR (qRT-PCR)

Total RNA was isolated from cultured cells using TRIzol reagent (Invitrogen, USA). 1.5µg purified RNA from each sample was reverse transcribed to the complementary DNA using HiScript III RT SuperMix for qPCR kit (Cat# 323, Vazyme Biotech, Nanjing, China). The cDNA was then used as template for quantitative PCR (qPCR), then were accomplished by using the ChamQ Universal SYBR qPCR Master Mix (Cat#Q711, Vazyme Biotech, Nanjing, China). The target genes BZLF1, BALF2, and BMRF1 expression levels were calculated using the 2^−ΔΔCt^ method. The relative levels of each sample were normalized to the β-actin housekeeping gene. Primer sequences are listed in Table 1.

**Table 1:**
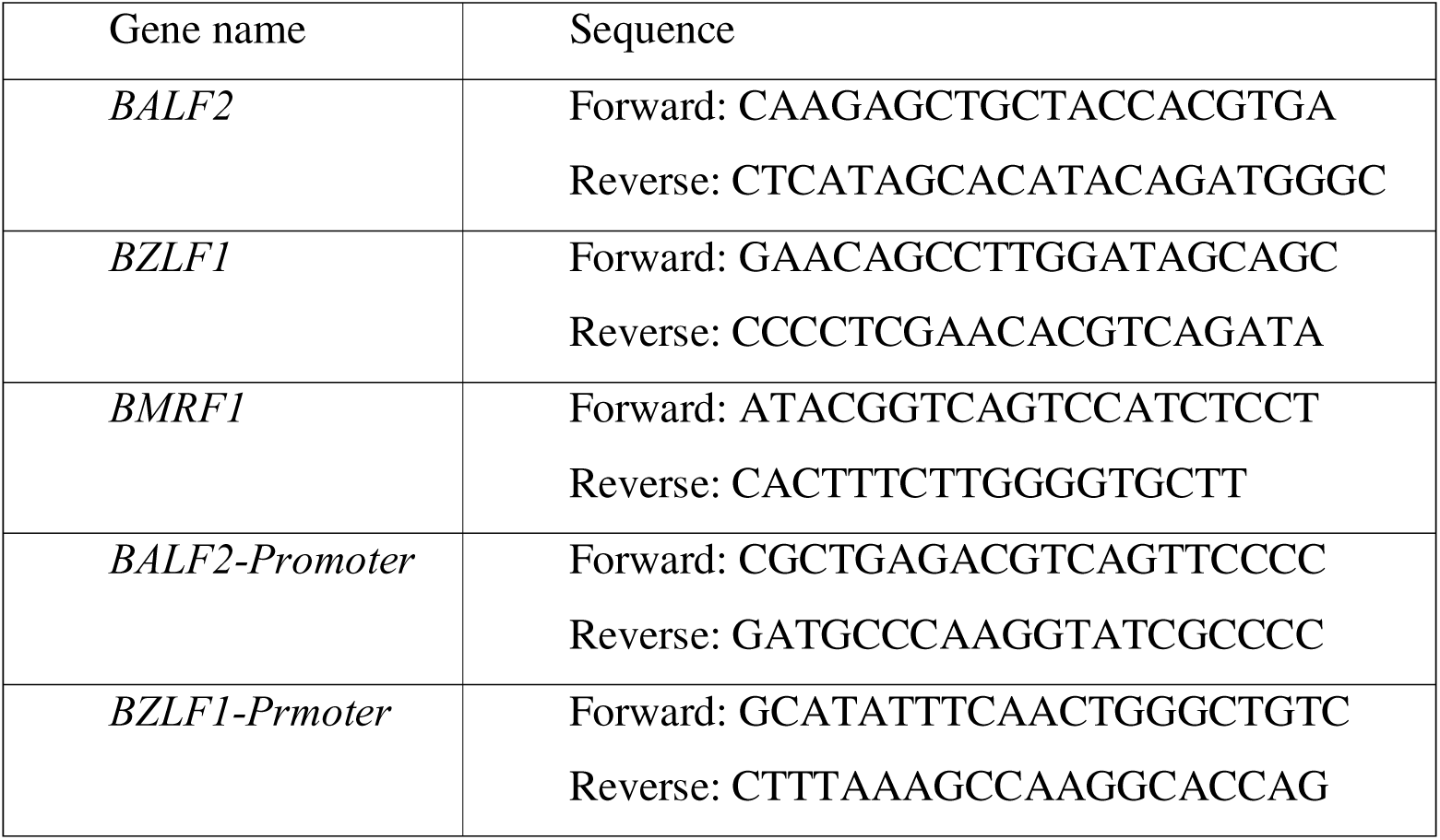
qRT-PCR primers used in this article: 5’-3’.

### 2.4 Electrophoretic mobility shift assay (EMSA)

The oligonucleotides correspond to SP1 binding sites in the BZLF1 and BALF2 promoter regions (outlined in Fig. S3). In selected oligonucleotides guanine residue(s) were replaced with 8-oxoGua during synthesis by Sangon Biotech (Shanghai, China) (Table 2). The oligonucleotides were then annealed with their complementary strands to form duplex probes. To perform the EMSA experiment, we utilized LightShift Chemiluminescent EMSA Kit (Cat# 20148, Thermo Scientific, USA). To examine the interaction of OGG1 with wild type DNA or 8-oxoGua DNA probe, 1 pmol Biotin-labeled duplex oligo was mixed with 10 ng OGG1 Recombinant protein in a total volume of 10 µL binding buffer including 10 mM of Tris-HCl (pH 7.5), 5 mM of NaCl, 1 mM of EDTA, 1 mM of DTT and 1 mg/mL BSA. The binding assay was performed for 10 minutes at 4[. The reaction mixtures were separated at 100 V for 90 minutes in a 6% non-denaturing polyacrylamide gel (0.5 × TBE) at 4°C.

**Table 2:**
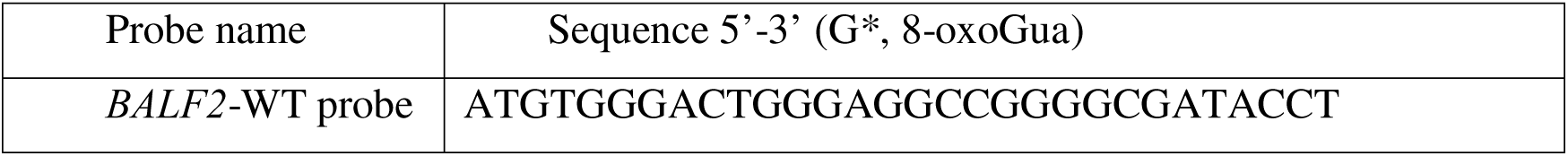
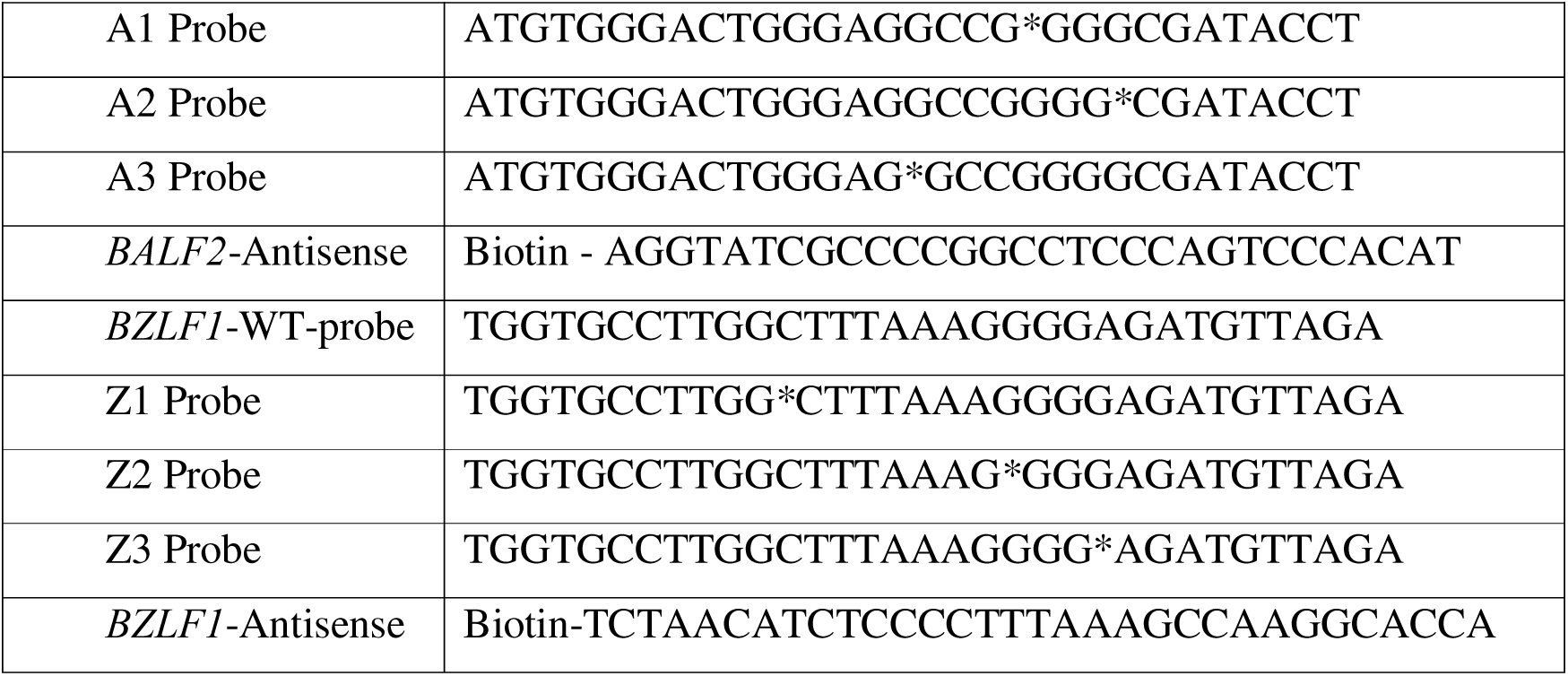
Oligos used in EMSA assay (5‘-3’)

### 2.5 Western blotting

Akata-EBV, C666-1 or B95.8 cells were treated as in 2.2. The obtained whole cell extracts were mixed with 2 × SDS sample buffers, separated by 10% of SDS-PAGE, and transferred onto nitrocellulose membranes. Membranes were blocked with 5% of skim milk (in 0.1% of TBST) for 1 hour and incubated with primary Abs overnight at 4°C. The dilution of monoclonal Abs against ZEBRA, Ea-D, ERK, p38, JNK or phosphorylation were 1:1000 and GAPDH was 1:8000. After three washes with TBST, membranes were incubated with secondary Abs and detected by ECL Plus western blot detection reagents.

### 2.6 Chromatin immunoprecipitation assay

2 × 10^7^ cells were cultured in 10-cm2 dish, pre-treated with or without 10 µM TH5487 for one hour, then, exposed to 20 ng/mL TPA plus 3 mM NaB for 20 hours. DNA-protein complexes were cross-linked with 1% of formaldehyde for 10 min at room temperature and sheared by sonication with 6-sec pulses (100-750 bp fragment) for six times (SCIENTA, Ningbo, China). DNA-protein complexes (10 µg of genomic DNA per sample) were immunoprecipitated using 2.5 µg of IgG or OGG1 Ab overnight at 4°C, then, incubated with 30 µL of magnetic beads for 2 hours. The precipitates were washed three times, de-cross-linked, and subjected to qPCR following the manufacturer’s instructions for the Simple ChIP® Enzymatic Chromatin IP Kit (Magnetic Beads) (Cat# 9003s, Cell Signaling Technology, USA). ChIP-qPCR data are presented as fold enrichment over IgG controls. Primer specificity and amplification efficiency were validated prior to analysis and is shown in Table 1. The fold enrichment of OGG1 on the specific region of EBV genome was calculated as described previously(22, 23).

### 2.7 Assessment of EBV copy number

B95.8 cells (5 × 105/ml) were mock or treated with individual OGG1 inhibitors (TH5487 or O8 -10 µM) for 1h, then TPA (20 ng/ml) +NaB (3 mM) was added. 24 h later cell suspensions were freeze (-80°C) thaw 3-times. Cell lysates were clarified by centrifugation (10,000 rpm for 10 min) and DNAs were extracted from supernatant fluids using virus DNA/RNA extraction kit (Cat# SDK60116, BioPerfectus, China). Equal amounts of DNA were used to determine copy numbers of EBV genome by EB virus nucleic acid detection kit (fluorescence PCR) following the instructions of manufacturer (JC60106N, BioPerfectus, China).

### 2.8 ROS assay

CM-H2DCFDA (Cat# C6827, Invitrogen, USA) was used to measure changes in TPA+NaB-induced ROS levels in Akata-EBV, C666-1 and B95.8 levels according to the manufacturer’s instruction(24). Briefly, cells were seeded on 12-well plates for 18 hours, then exposed to 20 ng/ml TPA and 3 mM NaB for various lengths of time and incubated with 5 μM CM-H2DCFDA at 37°C for 10 minutes. Cells were washed twice with Extracellular Solution (Cat# C0216, Beyotime Biotechnology, China) and lysed with 800μL RIPA (Cat# 89901, Thermo Fisher Scientific, USA) on ice, then obtain supernatant by centrifugation. DCF fluorescence in supernatant fluids were determined by ClarioStar microplate reader (BMG LABTECH, Germany) with excitation/emission at 495 nm/530 nm. Results are presented as fluorescence unit (FU) changes.

### 2.9 Statistical analysis

All experiments were performed at least three times. Statistical analyses were performed using Student’s t test. The data are presented as the mean ± the standard deviation. The level of significance was accepted at \**P* < 0.05, \*\**P* < 0.01, and \*\*\**P* < 0.001.

## 3. Results

### 3.1 Characterization of EBV reactivation across latency types I–III

Latent EBV can switch to lytic reactivation in response to ROS, with the magnitude and kinetics shaped by latency program, host cell context, and the triggering stimulus, but the mechanism is unclear(25–27). To characterize reactivation, we used Akata-EBV (latency I), C666-1 (latency II)(28), and B95.8 (latency III)(29, 30) cells as our experimental system. Cells were treated with TPA (20 ng/mL) and the histone deacetylase inhibitor sodium butyrate (NaB, 3 mM), a combination reported to elevate ROS(6, 27). To establish appropriate time points for detailed analysis, cell lysates were collected over time to measure EBV reactivation at mRNA and protein levels.

The pattern of lytic gene expression varied between latency types. In type I latency (Akata-EBV cells), TPA+NaB induced *BZLF1* (encoding the IE transcriptional activator Zta protein) after a 12 h incubation period and reached a maximum (>70-fold) ∼20 h post-treatment (Fig. 1A), indicating relatively delayed activation. In C666-1 (latency II), *BZLF1* increased rapidly (from 1 h), with oscillatory increases through 24 h (Fig. 1B). In contrast to Akata-EBV and C666-1, in B95.8 cells, TPA +NaB, *BZLF1* mRNA levels increased steadily, starting at 1 h post-treatment reaching peak expression between 20-24 hours (Fig. 1C). These patterns, together with the greater permissiveness of latency III, potentially reflect broader viral gene expression and possibly host transcriptional support.

**Figure 1.**
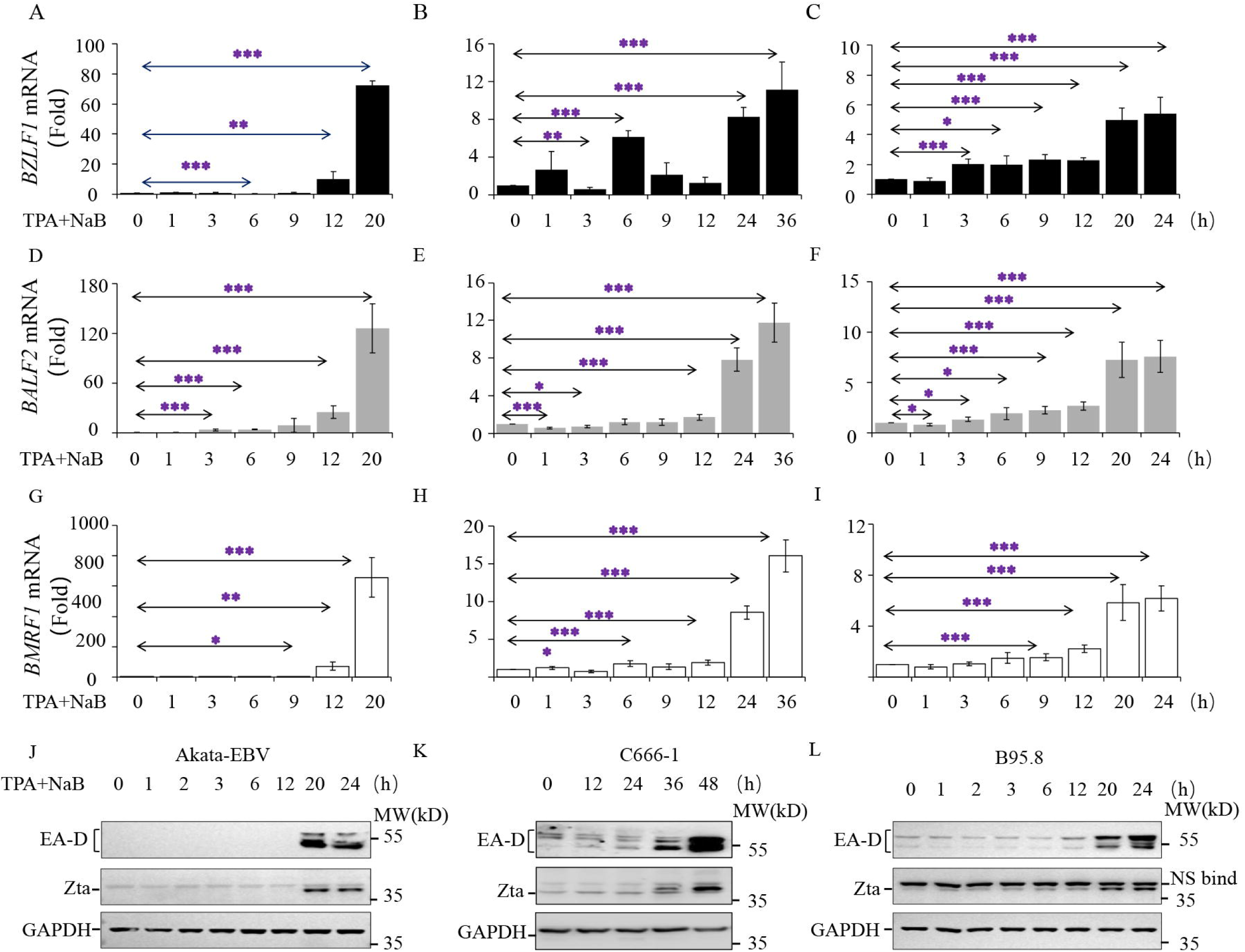
Kinetic expression of EBV IE and early genes at mRNA and protein levels in response to TPA+NaB. Cells were co-exposed to 20 ng/ml TPA and 3 mM NaB, and mRNA as well as protein levels encoded by (*BZLF1*, *BALF2* and *BMRF1*) were determined at time points indicated. The expression at mRNA levels of *BZLF1*, *BALF2*, *BMRF1* genes in Akata-EBV (**A, D, G**), C666-1 (**B, E, H**) and B95.8 (**C, F, I**) cells were determined by qRT-PCR. Panels **J** (Akata-EBV), **K** (C666-1) and **L** (B95.8) display the corresponding protein level as determined by Western blotting. All experiments were performed at least three times and data are presented as means ± the standard deviation. \*\*\**P* < .001, \*\**P* < .01, \**P* < .05.

Downstream early genes showed coordinated induction. In Akata-EBV, *BALF2* (ssDNA-binding protein) and *BMRF1* (DNA polymerase processivity factor) increased nearly in parallel with *BZLF1*. In C666-1 and B95.8, TPA+NaB overcame latency-associated silencing by ∼3 h, with *BALF2* and *BMRF1* cresting at 20-36 h, albeit with model-specific magnitudes (Fig. 1D-I).

Protein expression kinetics mirrored those of mRNA, albeit with latency-type dependent delays. Zta protein increased in Akata-EBV from ∼20 h (Fig. 1J), in C666-1 appeared at low levels by 12 h then rose strongly at 48-72 h (Fig. 1K), while in B95.8 Zta accumulated from ∼20 h (Fig. 1L). BMRF1 (EA-D) was tightly regulated in Akata-EBV but followed similar temporal patterns in C666-1 and B95.8 (Fig. 1J-L). Together, these data indicate TPA+NaB treatment overrides latency silencing, likely through TPA-mediated cell activation signaling and NaB-induced chromatin modification by inhibiting histone deacetylases, with the specific latency program shaping the timing and amplitude of lytic gene activation.

**3.2 Pharmacological inhibition of OGG1 suppresses TPA+NaB-induced EBV lytic gene expression**

Treatment with TPA+NaB induced significant increases intracellular ROS levels from 1h post-treatment (**Fig. 2A**) consistent with prior reports (6, 27), which remained elevated through experimental period (24 h) in all three models (**Fig. S1**). ROS- and TPA-signaling can drive the expression of a wide range of host gene expression through activation of transactivators including NF-κB, AP1, GATA, Sp1 and Nrf2/C-Ets1(31). Additionally, during EBV replication OGG1 expression is increased to offset the accumulation of oxidized DNA bases (e.g., 8-oxoguanine) and preserve integrity of viral genome, in addition to maintaining cell viability allowing efficient virus persistence and replication(13). Therefore, we determined its expression in latently types. Co-treatment of cells with TPA+NaB transiently and modestly changed OGG1 expression at RNA levels in Akata-EBV (**Fig. 2B**, left columns) and C666-1 cells (**Fig. 2B**, middle columns) at time investigated, while significant increases were observed in B95.8 cells (latency III) as a function time (**Fig. 2B**, right columns). These results imply that TPA+NaB and ROS signaling have latency- and cell type-dependent impact on OGG1 expressions.

**Figure 2.**
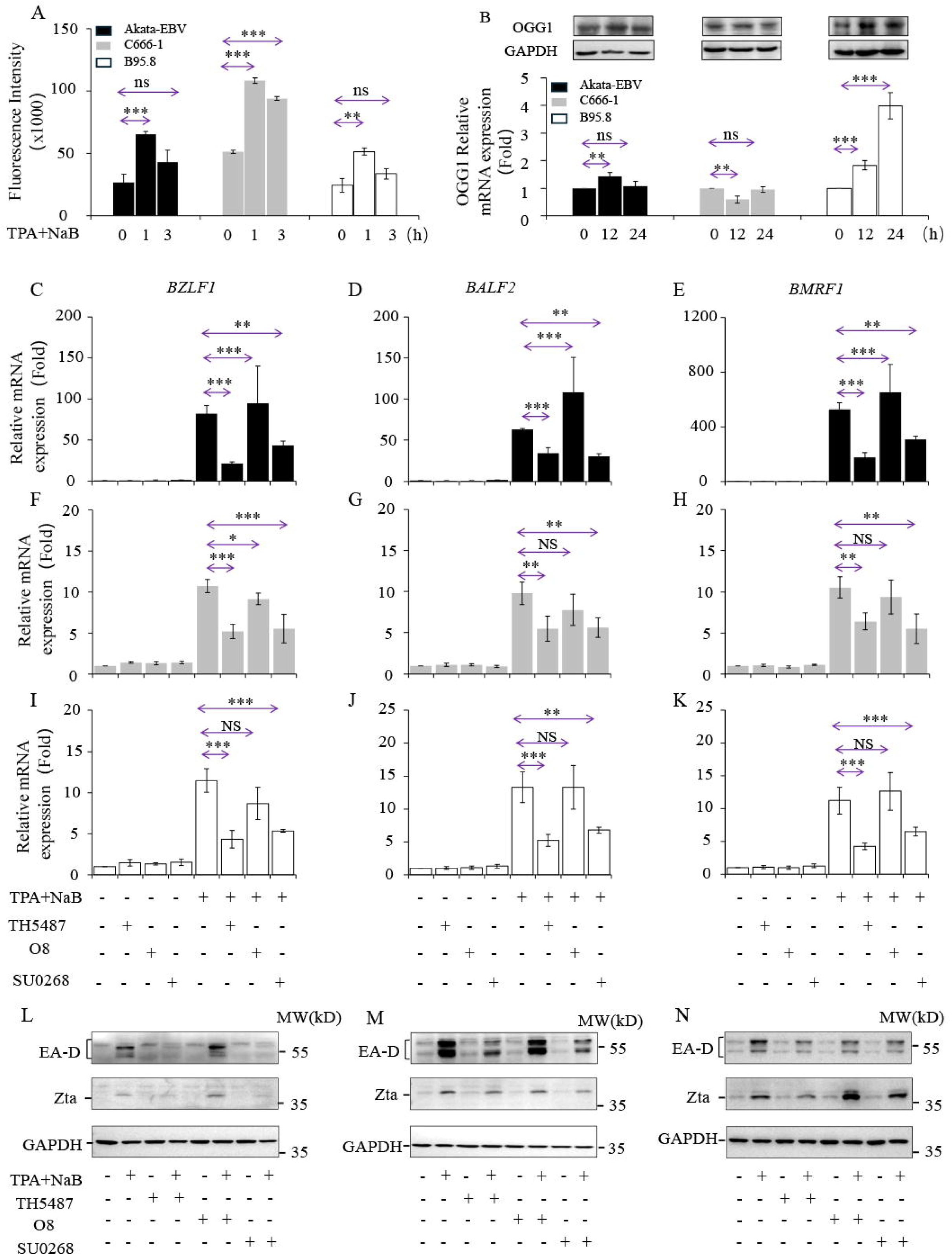
Inhibition of substrate recognition but not glycosylase activity of OGG1 decreases TPA+NaB-induced expression of EBV lytic genes. **A**, TPA+NaB addition of cells increase intracellular level of ROS as assessed by DH2CF-DA assays. **B** TPA+NaB-ROS alters OGG1 at mRNA (Akata-EBV (left), C666-1 (middle) and B95.8 (right), while there were no significant changes at protein levels. **C-K,** OGG1 binding but not its glycosylase activity is required for the expression of IE and early EBV genes. Cells were pre-treated with solvent or TH5487, SU0268 (10 μM) or O8 (10 μM), then activated by TPA+NaB (as in legend to Fig. 1). Cells were harvested at 20h post activation (see Fig. 1). TH5487 and SU0268 significantly decreases mRNA levels of BZLF1, BALF2 and BMRF1 and EBV protein levels (Zta, EA-D) in Akata-EBV (C, D, E and L), C666-1 (F, G, H and M) and B95.8 (I, J, K and N). O8 substantially increased mRNA levels in Akata-EBV (C-E), but no effect was observed in C666-1 (F-H) and B95.8 cells (I-K). O8 increased Zta and EA-D protein in all latency types. Changes in mRNA levels were determined by quantitative real-time PCR and protein level were determined by western blotting. Data were presented as averages ± SD from at least three independent experiments. \*\*\**P* < .001, \*\**P* < .01, \**P* < .05.

To investigate the role of OGG1 in EBV lytic gene expression, we initially considered genetic depletion strategies. However, nucleic acid insertion and genome editing technologies can induce cellular stress responses, thereby triggering EBV reactivation(27, 32). Consequently, we adopted pharmacological inhibition to examine the role of OGG1 in EBV reactivation. We employed well-characterized, structurally distinct small molecule inhibitors of OGG1, TH5487 and SU0268, which prevent OGG1 binding to intrahelical 8-oxoGua without inducing cytotoxicity(33, 34). Cells were pretreated for 1h with 10 μM inhibitor (optimized in preliminary studies), then exposed to TPA+NaB. In vehicle controls, TPA+NaB boosted *BZLF1*, *BALF2,* and *BMRF1* by ∼82-, 62-, and 529-fold, respectively, in Akata-EBV cells. Taking TPA+NaB induced increases as hundred percent, TH5487 pretreatment reduced *BZLF1, BALF2*, and *BMRF1* expressions by 79% ±7%; 48% ±12% and 75% ±14%, respectively (**Fig. 2C, D and E**). SU0268 produced comparable decreases (**Fig. 2C, D and E**). Consistent with the mRNA data, TH5487 and SUO268 lowered Zta and early antigen-diffuse (EA-D) protein levels (**Fig. 2L**). In C666-1, both inhibitors decreased mRNA levels of *BZLF1, BALF2*, and *BMRF1* significantly although extent of decrease averaged only by ∼40% (**Fig. 2F, G and H**). Decreases in Zta and EA-D protein were proportional with mRNA levels (**Fig. 2M**). In B95.8, where a small fraction of cells (∼1.5-2%) continuously entering the lytic cycle, the addition of TPA+NaB increased mRNA level of *BZLF1*, *BALF2*, and *BMRF1* by 11.5-, 13.2- and 11.2-fold, respectively compared to solvent only controls. TH5487 addition lowered mRNA levels of *BZLF1*, *BALF2*, and *BMRF1* by 61 ±9%; 67 ±13%, and 68 ±8%, respectively (**Fig. 2I, J, K**). The inhibitor SU0268 showing similar efficacy (**Fig. 2I, J, K**).

To examine, whether OGG1’s substrate binding and/or its enzymatic activity is required for activation of EBV lytic genes, we used O8, which inhibits OGG1 glycosylase activity but not DNA binding(35). Treatment with O8 did not decrease expression of *BZLF1*, *BALF2*, or *BMRF1* mRNAs in TPA+NaB-treated cells. In fact, O8 increased levels of Akata-EBV along with protein levels of Zta and EA-D substantially but not significantly (**Fig. 2C, D and E**). In latency II (C666-1) and III (B95.8) O8 had no effect on TPA+NaB-induced lytic gene expression (**Fig. 2C-K**). Because the OGG1 inhibitor O8 inhibits Schiff base formation during OGG1-mediated catalysis, these results support the possibility that specific binding of OGG1 to its DNA substrate rather than its glycosylase activity, promotes lytic gene transcription during reactivation. Importantly, TPA+NaB addition to B95.8 cells increased levels of EBV genome, which was decreased by TH5487 to pre-treatment levels (**Fig. S2**).

### 3.3 OGG1 does not lie upstream of MAPK activation during reactivation

Given that the OGG1-8-oxoG complex can function as a GEF to activate RAS and downstream MAPK or NF-κB signaling, thereby promoting gene expression(16, 17), we hypothesized that OGG1 might similarly promote EBV reactivation through protein phosphorylation-mediated signaling. Although EBV reactivation was observed in different latency models, subsequent mechanistic analyses were focused on Akata-EBV and B95.8 cells, which exhibited more robust and reproducible responses, to examine whether OGG1 acts upstream of TPA+NaB-induced signaling during the latent-to-lytic transition by evaluating the effects of OGG1 inhibitors on TPA+NaB-triggered MAPK, ERK and JNK signaling. Specifically, we assessed the phosphorylation of ERK1/2, JNK, and p38 MAPK in Akata-EBV and B95.8 cells, as these kinases are known to activate transacting factors that drive EBV lytic gene expression(9, 25, 36, 37).

In Akata-EBV cells, TPA+NaB triggered rapid ERK1/2 phosphorylation (Thr202/Tyr204) peaking at ∼1h and declining by 12h, modest early p38 phosphorylation (Thr180/Tyr182) with robust elevation at 20 h, and sustained JNK phosphorylation (Tyr185) across the time course (**Fig. 3A**). In B95.8, ERK1/2 phosphorylation always remained elevated, while p38 and JNK increased after ∼12 h (**Fig. 3C**).

**Figure 3.**
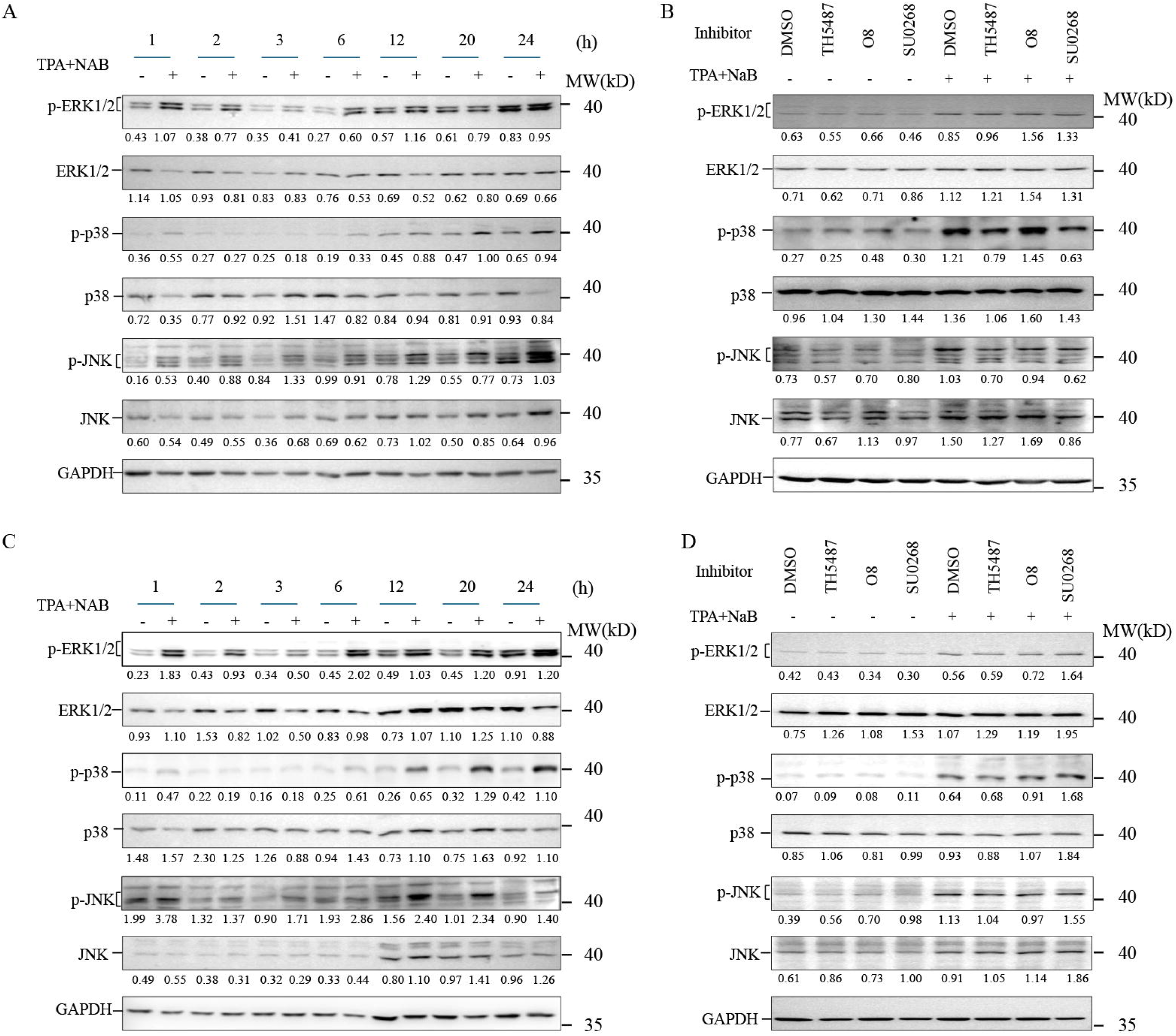
OGG1 does not impact TPA+NaB-induced ERK1/2, p38 and JNK phosphorylation. A, **C** Phosphorylation of ERK1/2 (Thr202/Tyr204), p38MAPK (Thr180/Tyr182) and JNK (Tyr185) after TPA+NaB treatment of Akata-EBV (A) and B95.8 (C) cells. Cells were activated by 20 ng/ml TPA and 3 mM NaB and cell lysates were prepared as indicated. **B, D** Lack of OGG1 inhibitor’s effect on the phosphorylation of ERK1/2 (Thr202/Tyr204), p38MAPK (Thr180/Tyr182) and JNK (Tyr185) in TPA and NaB treated Akata-EBV (B) and B95.8 (D) cells. Parallel groups of cultured cells were obtained OGG1 inhibitor TH5487, SU0268, O8 or solvent (DMSO) for 1h followed by addition of 20 ng/ml TPA and 3 mM NaB. Cell lysates were prepared at the indicated time points after activation to determine phosphorylation levels by Western blotting. Band intensities were quantified using ImageJ, and the relative protein expression levels were normalized to GAPDH.

Given that the most abundant phosphorylation of these kinases and expression of EBV IE and early genes are nearly parallel (**Fig. 1 and Fig. 3A, C**), we asked whether OGG1 inhibition by TH5487, SU0268 or O8 alters phosphorylation of MAPKs at key time points. Our results showed that pharmacological blockade of OGG1 binding (TH5487, SU0268), or inhibition of its glycosylase activity (O8) did not alter the phosphorylation of ERK1/2, p38 MAPK, or JNK at their respective peak times (**Fig. 3B, D**). These findings strongly suggest that OGG1’s contribution to lytic gene expression is largely independent of its GEF function to activate small GTPases and downstream MAPK pathway. Instead, these results are more consistent with a model through its association with oxidative DNA damage at viral regulatory regions and function as a transcriptional co-regulator at viral regulatory regions. Therefore, we next examined whether OGG1 is recruited to EBV lytic gene promoters during reactivation.

### 3.4 OGG1 is enriched at EBV lytic promoter regions during reactivation

To test whether TPA+NaB-induced ROS generates OGG1 substrate within the EBV genome, chromatin immunoprecipitation (ChIP)-coupled qPCR was performed (Materials and Methods). Primer pairs amplified *BZLF1* and *BALF2* promoter regions (nt 103,411–103,225 [187 bp], 164,771–164,922 [152 bp] aligned to EBV B95.8 genome V01555.2. Akata-EBV (type I latency) and B95.8 (type III latency) cells were analyzed.

In mock-treated Akata-EBV cells, OGG1 enrichment was 1.4-fold increase on *BZLF1* promoter compared with IgG control, but not on *BALF2* promoter. Pre-treatment with TH5487 for 1h had no significant effect under these conditions (**Fig. 4A**). TPA+NaB treatment of Akata-EBV cells significantly increased OGG1 enrichment on both *BZLF1* and BALF2 promoter sequences, which were decreased by TH5487 to pre-treatment levels (**Fig. 4A**). In mock-treated B95.8 cells, OGG1 enrichment (vs IgG) was ∼1.4-fold on *BZLF1* and 7.1-fold on *BALF2* sequences (**Fig. 4B**), suggesting that OGG1 may have a regulatory or repair function in latently infected cells (will be determined in future studies). Upon oxidative stress induced by addition of TPA+NaB, increased OGG1 enrichment over ∼2.5-fold and ∼3.2-fold (**Fig. 4B**). TH5487 prevented OGG1 enrichment to *BZLF1* and *BALF2* sequences in B95.8 cells (**Fig. 4A and B**). Since OGG1 binding is dependent on its DNA substrate, these data suggest that TPA+NaB-induced oxidative stress may promote generation of oxidative DNA lesions in EBV regulatory sequences and support inhibitor on-target activity at viral chromatin.

**Figure 4.**
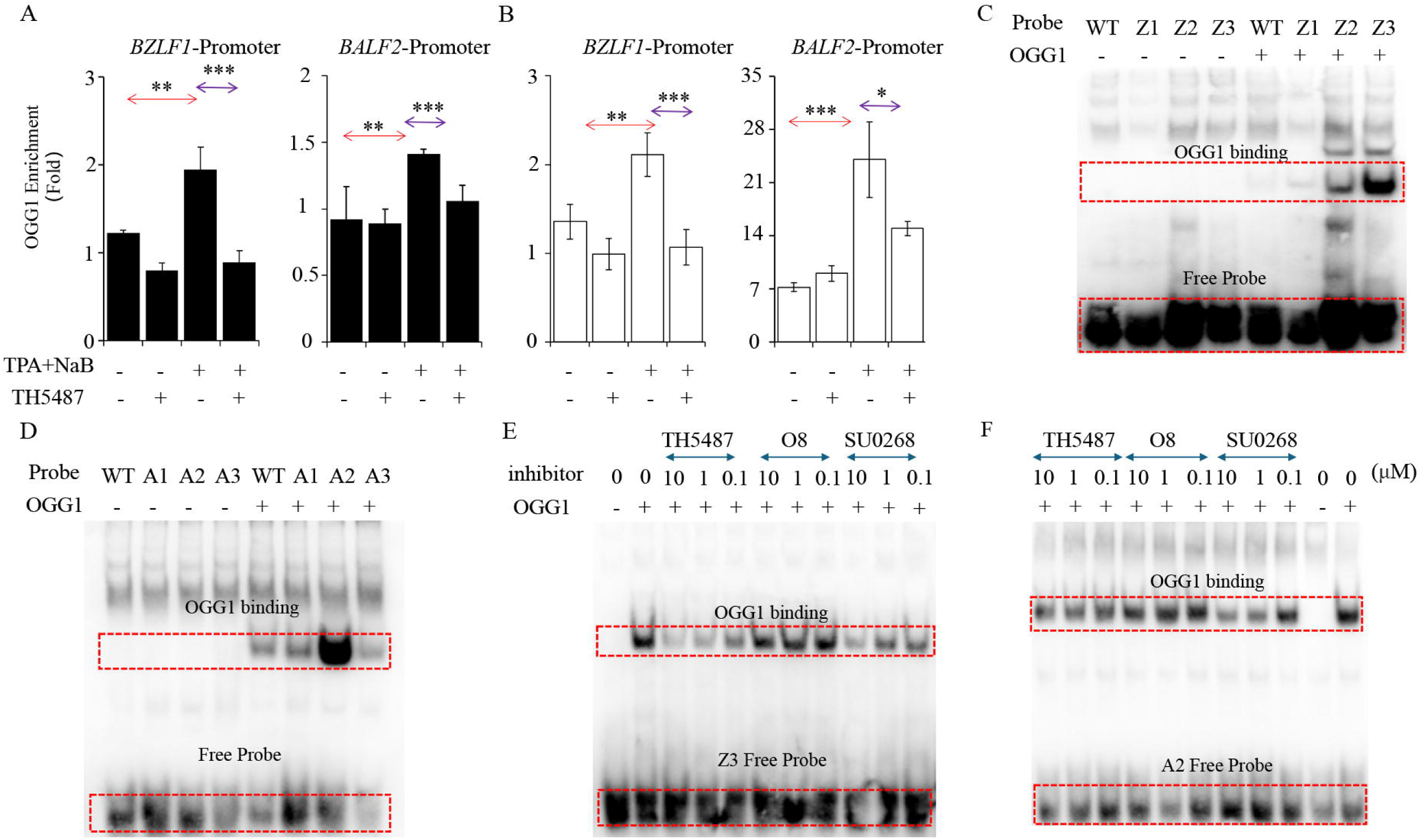
Enrichment of OGG1 at *BZLF1* and *BALF2* gene promoters. **(A** and **B)** Enrichment of OGG1 on *BZLF1* and *BALF2* promoter regions in Akata-EBV (A) and B95.8 (B) cells. Cells were pre-treated with 10 µM TH5487 for 1 h followed by 20 ng/ml TPA and 3 mM NaB and fixed at times of **max mRNA expression** (Fig. 1). ChIP was performed using an antibody to OGG1 or IgG (as a control). Enrichment of promoter regions were determined by qPCR. Fold changes were calculated as described in the Materials and Methods. (**C and D**) OGG1 preferentially binds to 8-oxoGua-containing DNA probes. Ten ng purified OGG1 was incubated with the indicated oligonucleotides on ice for 15□min. Samples were then subjected to a 6% non-denaturing gel electrophoresis at 4°C (Materials and Methods). **(E and F**) TH5487 and SU0268, but not O8 disrupt OGG1 binding to 8-oxoGua-containing probe. Ten ng OGG1 were incubated with the indicated concentrations of TH5487, SU0268 or O8 for 30 min at 37□ in 1x binding buffer, followed by the addition of probe for 15 min on ice. In C-F, the protein-DNA probe complexes were visualized using LightShift chemiluminescence EMSA kit. *\*\*\*P* < .001, *\*\*P* < .01, *\*P* < .05.

Next, we furthered these results, using *in vitro* studies --utilizing electrophoretic mobility shift assays (EMSA). EMSA was performed using DNA probes matching guanine-rich promoter segments including Sp1 sites (crucial for activation of the EBV BZLF1 and BALF2 promoters by Rta as they function as a multifunctional transcriptional hub to initiate lytic replication (38, 39). 8-oxoGua was placed synthetically, into sense strand at various distances from Sp1 binding elements and annealed with biotin-labeled complementary strand (Table 2). As shown in **Fig. 4C, D**, OGG1 binding was observed with 8-oxoGua-containing probes, both in the *BZLF1* (Z3) and *BALF2* (A2) probes in a DNA base composition manner and binding was especially strong when 8-oxoGua was placed at 3’-end of G series (A2 and Z3 probes). No interaction was observed between OGG1 and probe lacking 8-oxoGua.

To examine the specificity of OGG1-DNA probe interaction, we tested the effects of OGG1 inhibitors, TH5487, SU0268 and O8 (**Fig. 4E, F**). Ten µM TH5487 significantly inhibited OGG1 binding to the Z3 probe, while 1 µM TH5487 partially decreased binding. Similarly, SU0268 also decreased OGG1 binding to the Z3 probe (**Fig. 4E**). Additionally, both TH5487 and SU0268 prevented OGG1 enrichment on the A2 probe (**Fig. 4F**). In contrast, O8 (which inhibits the base excision activity of OGG1) did not disrupt OGG1-DNA binding; instead, OGG1 remained bound to 8-oxoGua-containing probe, resulting in increased accumulation of OGG1-DNA complexes (**Fig. 4E, F**). These orthogonal data support a model in which OGG1 is recruited to EBV promoters may be associated with oxidative DNA damage generated under TPA+NaB-induced conditions.

## 4. Discussion

EBV persists in host cells by establishing latency, during which the viral genome is chromatinized and tightly regulated by histone and DNA modifications, and transcriptionally silent except for a limited set of latency genes. Periodic reactivation into the lytic cycle is critical for viral propagation causing EBV-associated disease pathogenesis. Although diverse physical, chemical, and biological stressors are known to trigger reactivation, a unifying hallmark is the generation of ROS. Yet, the molecular mechanisms by which ROS bridge environmental stress to viral gene transcription remain incompletely defined(27, 40).

Here, we identified oxidative DNA lesion(s)-specifically 8-oxoGua and its cognate repair enzyme OGG1 as potential co-regulator(s) of EBV reactivation. Results showed that TPA(+NaB)-induced ROS was associated with increased OGG1 enrichment at *BZLF1* and *BALF2* promoter sequences. These findings extend prior observations of OGG1 functioning as a “DNA base modification reader” in host chromatin and demonstrate its involvement in regulating viral transcription. Importantly, pharmacological inhibition of OGG1-DNA binding suppressed lytic gene expression, highlighting its potential as a therapeutic target.

Treatment with TPA + NaB efficiently induced EBV lytic reactivation and ROS generation in Akata-EBV, C666-1, and B95.8 cells, accompanied by phosphorylation of ERK, p38 MAPK, and JNK, as reported previously(25, 26, 41). Although the OGG1-8-oxoGua complex has been shown to activate RAS-MAPK/NF-κB signaling, only when 8-oxoGua added exogenously(17), so OGG1 functioning as GEF can be excluded from activation processes of MAPK pathways in our system, indicating that OGG1 does not promote EBV reactivation through kinase-driven signaling cascades(17, 42). Instead, our data support a model in which OGG1 recognizes oxidized guanine and may associate with EBV lytic promoters to facilitate transcriptional activation.

Treatment with TPA + NaB significantly induced OGG1 expression only in B95.8 (latency III). This distinct increase in OGG1 expression may be linked to the expression of multiple viral latency genes, including EBV nuclear antigens (EBNA1/2/3A–3C/LP), latent membrane proteins (LMP1/2A/2B), EBV-encoded small RNAs (EBERs), and microRNAs (43). In latency I and II, OGG1 expression was transiently altered, likely not affected by signaling cascades and activation of trans-acting factors induced by TPA + NaB. Whether OGG1 contributes to latency maintenance or simply marks lytic readiness remains a key question for future studies. These data are in line with EBV infection-induce increases in the cellular antioxidant defense via the expression of DNA repair proteins, including OGG1, MTH1 the later enzyme hydrolyzes oxidized 8-oxo-2′-deoxyguanosine triphosphate to prevent its incorporation into viral (and host) DNA, and MUTYH (adenine/guanine-specific adenine DNA glycosylase), which excises adenine mis-incorporated opposite 8-oxoGua to ensure genome fidelity (13).

OGG1 binding to 8-oxoGua in regulatory sequences was further confirmed by EMSA, which showed OGG1 association with probe only in the presence of its substrate. Since OGG1 binding to DNA is dependent on the presence of substrate (e.g., 8-oxoGua or FapyGua (21, 44, 45), the ChIP and EMSA results support the possibility of generation of one of these substrates in the latent EBV genome due to ROS produced by TPA (+NaB) treatment independently from latency type.

To further explore this possibility, we utilized OGG1-specific inhibitors. We first used TH5487 and SU0268 to prevent OGG1 from binding on its intrahelical substrate(s) (33, 34). These inhibitors occupy the OGG1 active site and block its DNA binding, significantly decreased mRNA and protein levels of IE and early EBV genes. To confirm the specificity of these inhibitors, we also used O8, which inhibits the glycosylase activity of OGG1 without disrupting its DNA binding(35). O8 prolongs OGG1 association with intrahelical 8-oxoGua, thereby increasing DNA occupancy of host transacting factors (46, 47). Interestingly, O8 resulting in increased expression or had gene specific effect, which was evident only in latency I (Akata-EBV cells) or after primary infection of THP-1 cells. These data support the hypothesis that OGG1-substrate interactions in viral DNA may contribute to initiation of reactivation and expression of EBV IE and early genes.

Data from previous studies show that under oxidative stress, cysteine residues of OGG1 are oxidatively modified, resulting in decreased enzymatic activity, which leads to prolonged OGG1 engagement with its DNA substrate(22, 42). Based on this, we speculate that ROS generated by TPA (+NaB) treatment induces OGG1 substrates in G:C-rich promoter regions of IE and early genes, reduces OGG1 turnover on 8-oxoGua, and triggers site-specific allosteric DNA alterations(48). Indeed, transcription factor binding sites within EBV regulatory regions-including the BZLF1 promoter-are CpG-rich and guanine-rich including Sp-1 recognition sequences (49, 50), making 8-oxoGua formation and OGG1 recruitment plausible, as supported by ChIP analysis.

Combined with previous research, OGG1 not only participates in DNA repair but also facilitates transcription factor recruitment in chromatin to modulate gene expression(19, 22, 42, 51–53). Therefore, it can be proposed that OGG1-associated viral gene expression may result from local allosteric changes in EBV promoters at 8-oxoGua-adjacent sequences that promote Zta binding to viral DNA in a manner mechanistically analogous to host transcription factor binding (e.g., NF-κB, AP1, MYC, SP1/3, SMAD, HIF-1) in chromatinized DNA(22, 42, 53, 54). This is supported by emerging data suggesting that OGG1 binding to gene regulatory sequences exerts an epigenetic role for 8-oxoGua(21, 55).

The EBV genome in virions lacks epigenetic marks, allowing immediate lytic gene expression upon primary infection (4, 54). During latency, however, chromatinization and CpG methylation suppress lytic promoters, a process mediated by host DNA methyltransferases(4, 56). Emerging evidence also highlights OGG1’s role in epigenetic regulation through modulation of CpG methylation(19, 57). Specifically, OGG1 bound to 8-oxoGua in chromatinized DNA recruits Ten-eleven translocations (TET), which can convert methylated cytosine (5mC) to 5-hydroxymethylcytosine (5hmC) (58). TET enzymes further oxidize 5hmC into 5-formylcytosine and 5-carboxylcytosine, which is excised by thymine DNA glycosylase and the gap repaired by DNA BER (59) resulting in removal of the suppressing mark. Moreover, 8-oxoGua itself in the chromatin decreases the binding affinity of DNA methyltransferases and OGG1 has increased 8-oxoGua binding within or in proximity of methylated CpG sites. Thus, OGG1 can significantly interfere with function/expression of latency-associated proteins while also acting as a transcriptional coregulator of lytic genes(53). These observations suggest a dual role for OGG1 in regulating both latency and lytic gene transcription. Further investigation of these interactions will elucidate novel molecular mechanisms underlying this process.

A unifying feature of EBV reactivation is ROS generation. This study identifies the oxidative DNA base lesions 8-oxoGua-recently recognized as epigenetic-like marks in EBV genome in latently infected cells. OGG1, the DNA glycosylase responsible for 8-oxoGua repair, may also function as an epigenetic-like reader and modulator of gene expression under oxidative stress conditions. This study provides initial evidence that OGG1-8-oxoGua interactions may contribute to regulation of the EBV lytic cycle from latency. Importantly, pharmacological inhibition of OGG1, particularly approaches that prevent DNA binding, may represent a potential strategy for reducing EBV reactivation in EBV-associated diseases.

Several limitations of this study should be acknowledged. First, although our findings consistently demonstrate that pharmacological inhibition of OGG1 significantly suppresses EBV lytic gene expression, the precise distribution and dynamics of oxidative DNA lesions across the EBV genome remain unclear. Moreover, a direct causal relationship among oxidative DNA damage, OGG1 recruitment to viral regulatory regions, and promoter-specific transcriptional activation of EBV lytic genes has not yet been fully established. Second, OGG1 may facilitate EBV lytic reactivation through interactions with specific transcription factors, chromatin remodeling complexes, or other transcriptional regulatory machinery; however, the downstream transcriptional regulatory network and critical molecular partners involved in this process remain to be further characterized. Third, stable genetic manipulation approaches, including siRNA-mediated knockdown and CRISPR/Cas9-mediated knockout of OGG1, remain technically challenging in EBV latency models because nucleic acid delivery itself can induce cellular stress responses and nonspecific EBV reactivation, thereby complicating interpretation of the results. In addition, further chromatin-level investigations, including promoter-specific oxidative DNA damage mapping, OGG1 occupancy analysis, and ChIP-based transcriptional studies, will be necessary to more precisely define the mechanistic role of OGG1 in EBV latency disruption. Finally, the present study was primarily conducted in established EBV-positive cell models, and additional studies using primary cells and in vivo systems will be important to validate the physiological relevance of these findings. Despite these limitations, our study provides new evidence supporting a noncanonical role of OGG1 in oxidative stress-associated EBV lytic reactivation and highlights OGG1 as a potential therapeutic target for controlling EBV reactivation and EBV-associated diseases.

## Supporting information

Supplementary_Material.docx

## 6. Author Contributions

**Wenjing Hao**

Contributed equally to this work with: Wenjing Hao, Xiaoqi Wang

ROLES: Conceptualization, Data curation, Formal analysis, Funding acquisition, Investigation, Methodology, Validation, Writing – original draft, Writing – editing

**Xiaoqi Wang**

Contributed equally to this work with: Wenjing Hao, Xiaoqi Wang

ROLES: Data curation, Funding acquisition, Formal analysis, Validation, Writing – review & editing

**Jiabo Li**

ROLES: Validation

**Lang Pan**

ROLES: Conceptualization, Writing – review & editing

**Xueqing Ba**

ROLES: Conceptualization, Writing – review & editing

**Gang Liu**

ROLES: Writing – review & editing

**Istvan Boldogh**

ROLES: Conceptualization, Supervision, Writing – editing manuscript

**Ziyuan Duan**

ROLES: Conceptualization, Funding acquisition, Investigation, Project administration, Resources, Writing – original draft, Writing – review & editing

## 8. Acknowledgments

This is a short text to acknowledge the contributions of specific colleagues, institutions, or agencies that aided the efforts of the authors.

