## Supplementary_Material.docx for "OGG1-Binding to Oxidized Guanine Base in Viral DNA Overcomes Epstein–Barr Virus Latency"

Preprint version: May 2026.

*** Correspondence:**
Corresponding Author: Ziyuan Duan


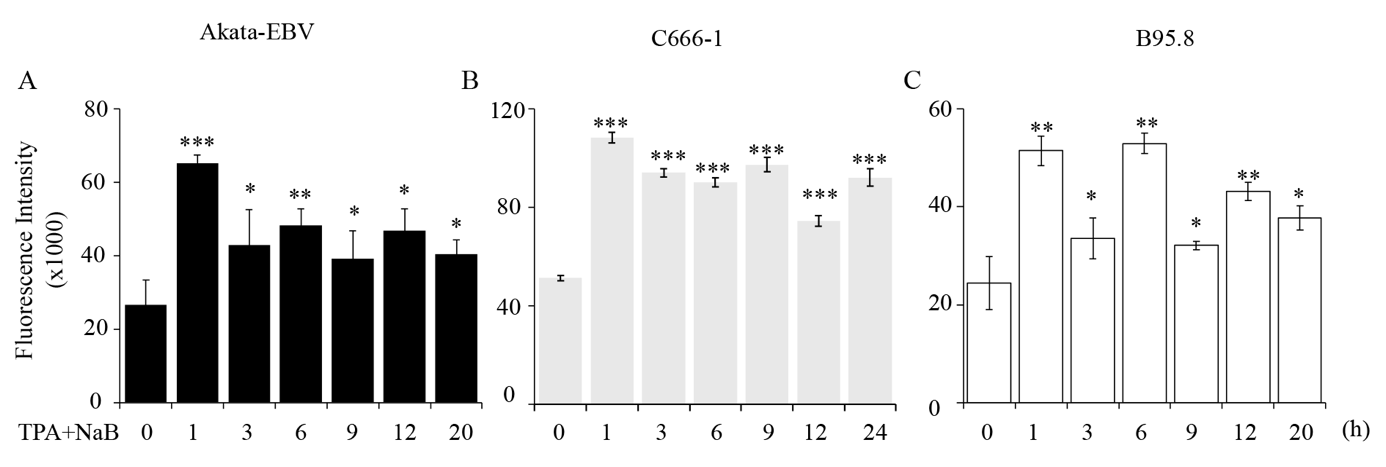

**Figure S1. TPA+NaB addition to cells increase intracellular level of ROS as assessed by DH2CF-DA assays.** Cells were co-exposed to 20 ng/ml TPA and 3 mM NaB, and intracellular level of ROS levels were determined at time points indicated.

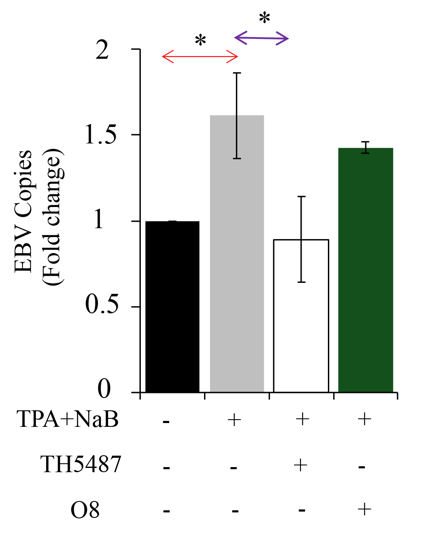

**Figure S2.** Inhibiting OGG1 substrate recognition suppresses EBV lytic replication. B95.8 cells were pre-incubated with TH5487 (10 μM) or O8 (10 μM) for 1h, followed by addition of TPA (20 ng/mL) and NaB (3 mM) for 24 h. EBV DNA copy numbers were quantified by qPCR.

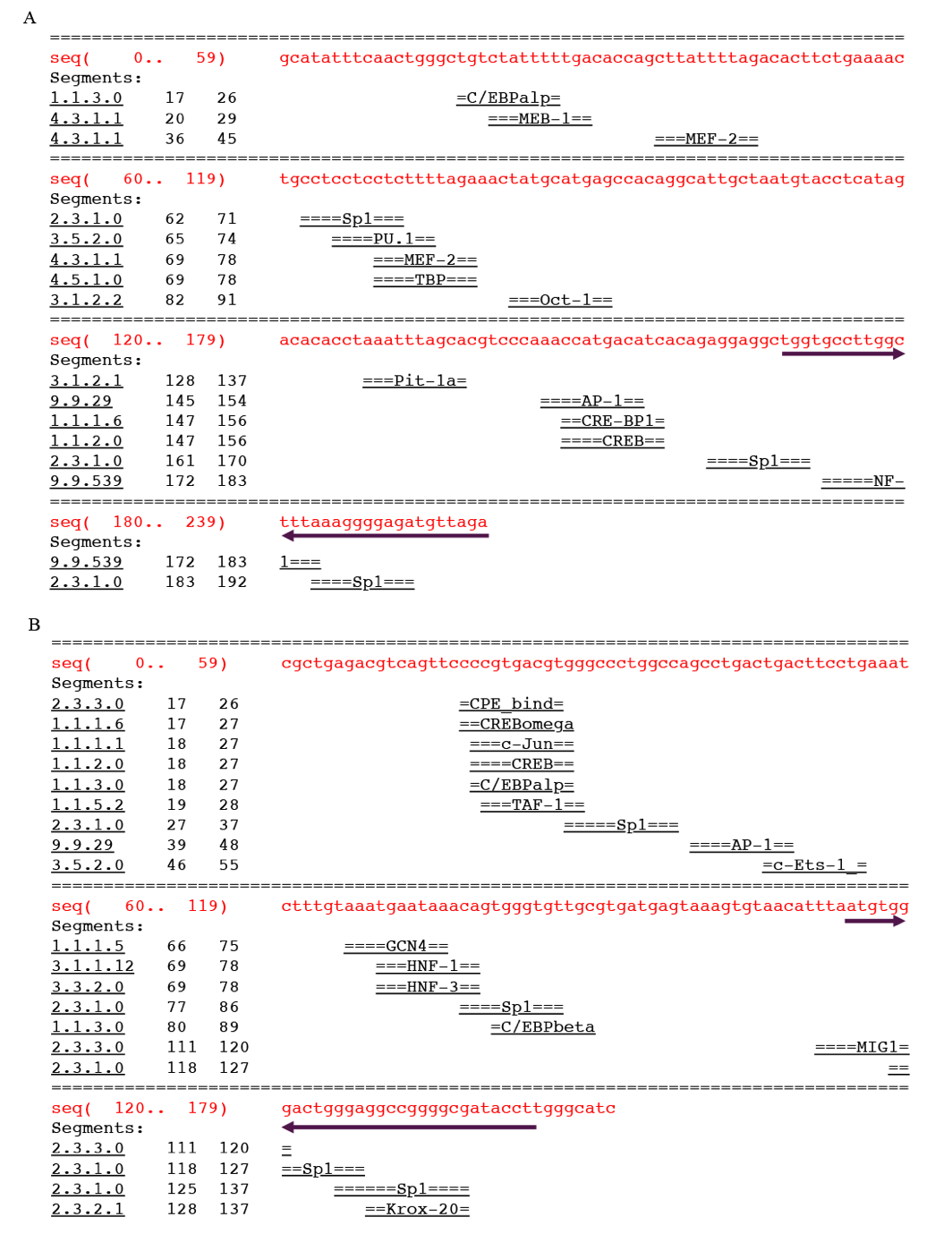

**Figure S3. Prediction of transcription factor binding sites in the EBV BZLF1 and BALF2 promoter regions.** (A) Schematic representation of the predicted transcription factor binding sites within the *BZLF1* promoter sequence. The putative binding motifs for transcription factors (including C/EBPβ, NF-κB, AP-1, Oct-1 and Sp1) are marked by horizontal bars below the corresponding regions, as identified using the AliBaba2.1 program based on the TRANSFAC database. (B) Predicted transcription factor binding sites within the *BALF2* promoter region. The prediction was performed using the AliBaba2.1 program based on the TRANSFAC database with the following parameters: Pair similarity to known sites = 64, matrix width = 10 bp, minimum number of sites = 4, minimum matrix conservation = 75%, sequence-to-matrix similarity threshold = 1%, and factor class level = 4. Purple arrows indicate the locations of EMSA probe regions used in this study. (<https://gene-regulation.com/pub/programs/alibaba2/index.html>)
